# Kynurenine aminotransferase II inhibition promotes sleep and rescues impairments induced by neurodevelopmental insult

**DOI:** 10.1101/2022.09.20.508758

**Authors:** Snezana Milosavljevic, Andrew K. Smith, Courtney J. Wright, Homayoun Valafar, Ana Pocivavsek

## Abstract

Dysregulated sleep is commonly reported in individuals with neuropsychiatric disorders, including schizophrenia (SCZ) and bipolar disorder (BPD). Physiology and pathogenesis of these disorders points to aberrant metabolism, during neurodevelopment and adulthood, of tryptophan via the kynurenine pathway (KP). Kynurenic acid (KYNA), a neuroactive KP metabolite derived from its precursor kynurenine by kynurenine aminotransferase II (KAT II), is increased in the brains of individuals with SCZ and BPD. We hypothesize that elevated KYNA, an inhibitor of glutamatergic and cholinergic neurotransmission, contributes to sleep dysfunction. Employing the embryonic kynurenine (EKyn) paradigm to elevate KYNA in the fetal brain, we presently examined whether reducing KYNA in adulthood by pharmacologically inhibiting KAT II would improve sleep quality. Pregnant Wistar rats were fed either kynurenine (100 mg/day) (EKyn) or control wet mash (ECon) from embryonic day (ED) 15 to ED 22. In adulthood, male and female offspring were implanted with devices to record electroencephalogram (EEG) and electromyogram (EMG) telemetrically for continuous sleep-wake data acquisition. Each subject was treated with either vehicle or PF-04859989 (30 mg/kg, s.c.), an irreversible KAT II inhibitor, at zeitgeber time (ZT) 0 or ZT 12. KAT II inhibitor improved sleep architecture maintaining entrainment of the light-dark cycle; ZT 0 treatment with PF-04859989 induced transient improvements in rapid eye movement (REM) and non-REM (NREM) during the immediate light phase, while the impact of ZT 12 treatment was delayed until the subsequent light phase. PF-04859989 administration at ZT 0 enhanced NREM delta spectral power and reduced activity and body temperature. In conclusion, reducing de novo KYNA production alleviated sleep disturbances and increased sleep quality in EKyn, while also improving sleep outcomes in ECon offspring. Our findings place attention on KAT II inhibition as a novel mechanistic approach to treating disrupted sleep behavior with potential translational implications for patients with neurodevelopmental and neuropsychiatric disorders.

## Introduction

More than one third of the global population currently reports problems with sleep (1). In patients with severe psychiatric disorders, the risk of irregular sleep patterns is two-fold higher compared to healthy controls, and thus, expectedly, the prevalence of sleep disturbances is 80% amongst individuals with schizophrenia (SCZ) and bipolar disorder (BPD) (2, 3). The causation and relationship between psychiatric disorders and sleep is complex and bi-directional (4, 5). As sleep is a universal, physiological process that serves a restorative function to the body, the loss and disruption of sleep in patients with psychiatric illness often exacerbates their symptom severity. For decades, besides benzodiazepines and melatonin (6, 7), clinical therapies for sleep disorders are associated with increased incidence of adverse effects such as sedation, daytime somnolence, and cognitive decline (8, 9). Therefore, optimizing novel therapies for clinical treatment of sleep disorders related to neuropsychiatric conditions has emerged as a high priority to improve health outcomes for patients.

Metabolites of the kynurenine pathway (KP) play a major role in both physiology and pathogenesis of major psychiatric disorders (10, 11). More than 95% of the ingested essential amino acid tryptophan is catabolized by the KP to generate nicotinamide adenine dinucleotide (NAD^+^), the ubiquitous co-factor and crucial energy source, as well as several neuroactive metabolites (12, 13). Elevated levels of kynurenic acid (KYNA), a kynurenine-derived gliotransmitter that inhibits glutamatergic and cholinergic neurotransmission (14–16), have been found in the postmortem brain and cerebrospinal fluid of individuals with SCZ and BPD (17–21). Inhibition of kynurenine aminotransferase (KAT) II, an enzyme which accounts for ~75% of the brain KYNA neosynthesis (22), elicits reduction of KYNA in the brain (23–25), which could have a direct impact on brain function and neurobiological processes relevant to SCZ and BPD.

Elevations in KYNA have also been implicated in pathogenic mechanisms that contribute to aberrant neurodevelopment which precedes the clinical onset of SCZ and BPD (26–28). Embryonic kynurenine (EKyn) experimental paradigm was established as a neurodevelopmental model that simulated in utero upregulation of the KP (27, 29). EKyn pregnant rats are treated with kynurenine during the last week of rodent gestation, translating to the second trimester in human pregnancy (30, 31), to induce a prenatal increase in KYNA. Notably, young adult EKyn offspring demonstrate sex-specific and circadian-dependent changes in KP metabolism in the hippocampus and the prefrontal cortex (PFC)(29, 32–34), namely increased KYNA within the brain of male offspring(32, 35). Stark cognitive behavioral impairments have been reported across studies in EKyn offspring (29, 33, 36, 37), which may be related to notable disruptions in sleep architecture, including shorter rapid eye movement (REM) duration in males, and prolonged episodes of quiescent wake behavior in females (32, 38).

Our present goal was to pharmacologically inhibit KAT II in an effort to reduce brain KYNA synthesis and to promote healthier sleep dynamics in ECon and EKyn rats of both sexes. Subjects were treated acutely with PF-04859989 (25), a brain-penetrable and irreversible KAT II inhibitor, at the start of the light phase or dark phase. We determined that this compound enhanced REM and non-REM (NREM) sleep, and importantly, KAT II inhibition in EKyn offspring restored sleep physiology to the level of counterpart controls. Together, these findings place novel attention on the translational value of KAT II inhibition for treating sleep deficiencies and improving sleep architecture.

## Methods

### Animals

Pregnant Wistar rats, embryonic day (ED) 2, were obtained from Charles River Laboratories (Raleigh, NC, USA). All experimental animals were housed in a temperature- and humidity-controlled facility, fully accredited by the Association for Assessment and Accreditation of Laboratory Animal Care (AAALAC) at the University of South Carolina School of Medicine. The rats had free access to food and water and were maintained on a 12h/12h light-dark cycle, where lights on corresponded to zeitgeber time (ZT) 0 and lights off to ZT 12. All experimental protocols were approved by the Institutional Animal Care and Use Committee (IACUC).

### Embryonic Kynurenine (EKyn) Treatment

Rodent chow was finely ground in a blender, and each dam received approximately 30 grams of wet rodent mash daily (ECon). Embryonic kynurenine (EKyn) dams were fed 30 grams of rodent mash daily with 100 mg/day of L-kynurenine sulfate (purity 99.4%, Sai Advantium, Hyderabad, India), from ED 15 through ED 22 as previously described (29). From the day of birth, postnatal day (PD) 0, standard rodent chow pellets were provided to all animals ad libitum. On PD 21, male and female offspring were weaned and pair-housed by litter and sex. Offspring were used experimentally from PD 56 to PD 85 (experimental timeline in **Figure 1**). To minimize the contribution of individual prenatal litters, the distribution of progeny from any given litter was one to two pups per sex for all experiments.

**Figure 1.**
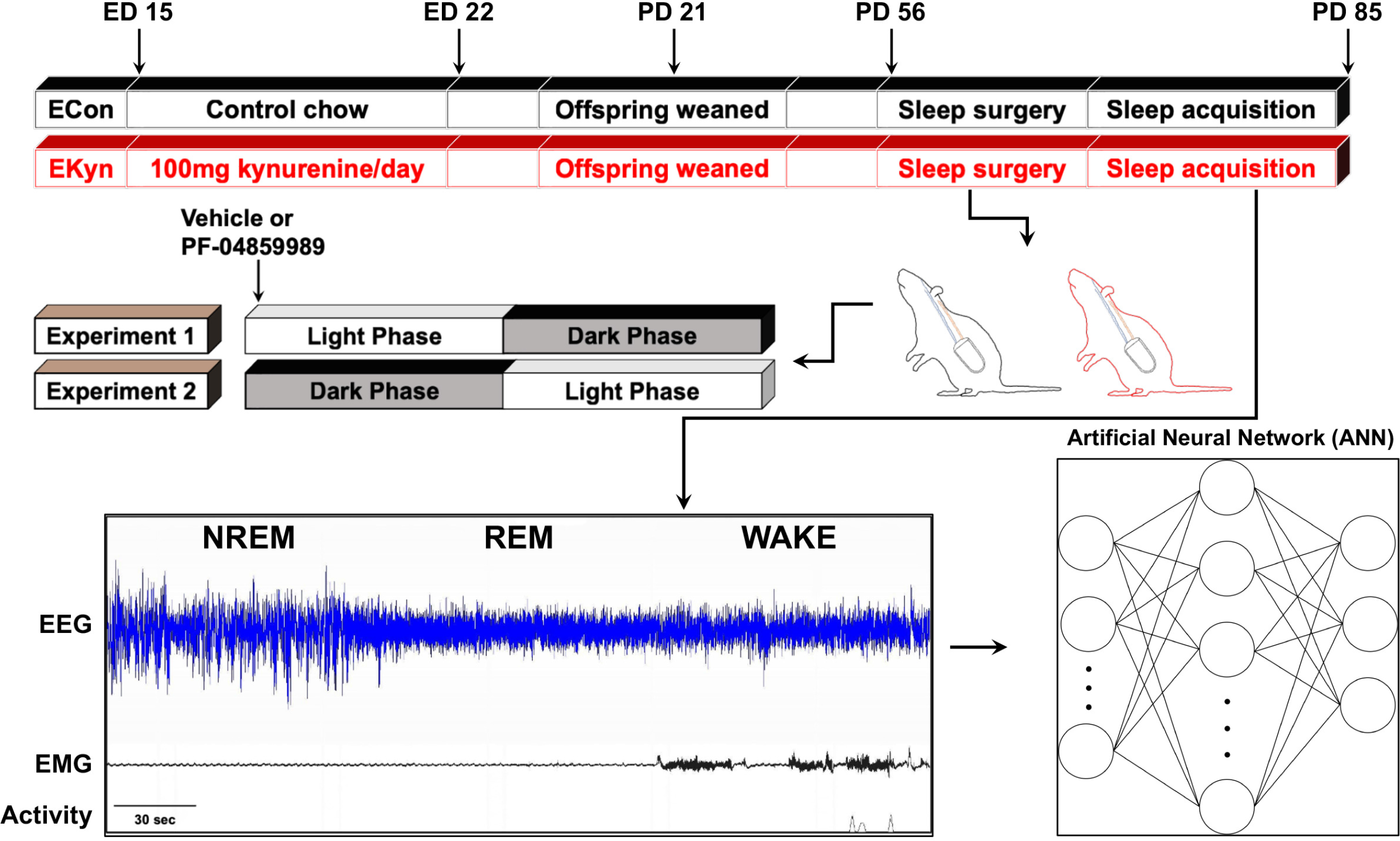
Schematic representation of experimental paradigm. Embryonic kynurenine (EKyn) treatment: from embryonic day (ED) 15 through ED 22 dams were fed daily with control diet (ECon) or diet laced with 100 mg kynurenine (EKyn). Sleep studies were performed in adult offspring between PD 56-85. In experiment 1, each subject was treated with vehicle or PF-04859989 (30 mg/kg) at the beginning of light phase, Zeitgeber time (ZT) 0. In experiment 2, each subject was treated with vehicle or PF-04859989 (30 mg/kg) at the beginning of dark phase, ZT 12. Electroencephalogram (EEG), electromyograph (EMG), and cage activity data were inspected in 10-second epochs and classified into one of three states: rapid eye movement (REM), non-REM (NREM), and wake. Artificial neural network (ANN) was implemented for highly accurate predictive classification of vigilance states.

### Surgery

Surgical implantation of telemetry transmitters (HD-S02; Data Sciences International (DSI), St. Paul, MN, USA) was according to the previously published protocols (38, 39). Briefly, animals were anesthetized using isoflurane anesthesia (induction 5%; maintenance 1.5 – 2.5%) and placed in a stereotaxic frame (Stoelting Co., Wood Dale, IL, USA) to secure the head. At the beginning of the surgical procedure, carprofen (5 mg/kg, s.c.) was administered as an analgesic. The telemetry transmitter was implanted intraperitoneally through a longitudinal incision made on the dorsal abdominal surface. Along the midline of the head and neck, another longitudinal incision was made to expose the skull and neck muscle, and electroencephalograph (EEG) and electromyograph (EMG) electrode leads were threaded subcutaneously to the incision. Two burr holes (0.5 mm diameter) were drilled and two surgical stainless-steel screws (P1 Technologies, Roanoke, VA, USA) were implanted at 2.0 mm anterior/ +1.5 mm lateral and 7.0 mm posterior/ −1.5 mm lateral relative to bregma. The two EEG leads were wrapped around the screws and secured with dental cement (Stoelting Co., Wood Dale, IL, USA). The two EMG leads were inserted directly into the dorsal cervical neck muscle approximately 1.0 mm apart and sutured into place. The dorsal incision was stapled closed with wound clips, and the skin along the head was sutured. Animals were allowed to recover postoperatively for at least 7 days prior to the start of experiments.

### Sleep Data Acquisition and Analysis

Sleep behavior was evaluated in a within animal design. Each subject received injections at ZT 0 for experiment 1 and at ZT 12 for experiment 2. For both experiments, each animal was treated with vehicle (ultrapure water) on the first data acquisition day and PF-04859989 (30 mg/kg, s.c.) during the other acquisition day (**Figure 1**). In an effort to match estrous phase in female subjects and allow ample time for drug wash out, treatments were spaced 3-4 days apart. Sleep data were collected for 24 hours after each injection. Total number of subjects recorded were N = 7-8 per group.

All sleep data were recorded in a designated room where animals were singly housed, and cages were placed on receivers that relayed data continuously to Ponemah 6.10 (DSI). Digitized signal data were processed offline with NeuroScore 3.0 (DSI). According to visual inspection of EEG/EMG waveforms and confirmation with a trained artificial neural network (ANN)(to 95% agreement; details below), 10-second epochs were classified into one of three states: wake, NREM, or REM. Wake was characterized by low-amplitude, high-frequency EEG combined with high-amplitude EMG; NREM by high-amplitude, low-frequency EEG combined with low-amplitude EMG; and REM by low-amplitude, high-frequency EEG combined with very low EMG tone and muscle atonia (see **Figure 1**). An uninterrupted episode of a single vigilance state lasting for at least one full epoch was defined as a bout, and a transition was scored when two or more epochs were noted as the new vigilance state.

The scored data for each vigilance state were analyzed in 1-hour time bins and 6-hour time bins for total duration, number of bouts, average bout duration, relative cage activity, and core body temperature. NREM and REM sleep onset and the number of transitions between vigilance states were determined for each 12-hour cycle. Power spectrum analysis during REM and NREM sleep, separated by phase, was evaluated with Discrete Fourier Transformation (DFT) to obtain the power spectrum for the following frequency bandwidths: delta (0.5–4 Hz), theta (4–8 Hz), alpha (8–12 Hz), sigma (12–16 Hz), and beta (16–20 Hz).

### Artificial Neural Network (ANN)

ANN is a supervised predictive machine learning technique that is popular due to its success in recognizing patterns. We have shown that uncalibrated ANN is highly accurate in classifying vigilance state (wake, NREM, or REM) in rats based on features extracted from their polysomnography (40). In brief, we presently calibrated and verified correctness of the ANN by using over 400 hours of continuously monitored EEG and EMG (scored by a domain expert as described above), and relative home cage activity. Signals were partitioned into 10-second epochs where each epoch was then individually transformed to the frequency domain using DFT. Power spectrum and total power were computed for 40 channels of equal width from 0 to 20Hz, resulting in the first 40 input features, the 41st feature was the average of the EMG signal for each epoch, and the 42nd feature was cage activity, derived parameter from Ponemah. Using a simple moving window (overlapping each neighboring window by four epochs) of five epochs long (constituting 50 seconds of raw signal), input samples were created to consist of 210 features (42 discrete features for each epoch in the window). For the reformatted input samples, we computed labels by taking the most common class from the five epochs in the window. To enforce generalization and eliminate memorization, we partitioned 64% of the data into training data, 16% into validation, and 20% into testing. We implemented a fully-connected shallow ANN with 210 input neurons, 512 hidden layer neurons, and 3 output neurons. The REctified Linear Unit (RELU) activation function was used in the hidden layer to introduce non-linearity. The three neurons in the output layer mapped to the three discrete vigilance states (wake, NREM, REM) with the Softmax activation function. Calibration of the network was performed on the training and validation data and continued until a sufficient plateau of the validation loss curve occurred. After training, we evaluated the ANN on testing data (data it has never seen before) using accuracy, F1 score, and a confusion matrix. We further scrutinized the ANN by visual inspection of its predictions of the testing data by a domain expert.

### Open Field Test

Open field test was used to examine exploratory behavior and spontaneous locomotor activity in rats. Experiments were conducted during the light phase (ZT 2 – 4) with lights on or during the dark phase (ZT 22 – 24) under red light. An open field arena (70cm × 70cm) was divided into a square central zone in the middle (35cm × 35cm), a peripheral zone which consisted of four corners (17.5cm × 17.5cm), and a perimeter zone (17.5cm from arena walls). Each rat was placed in the center of the open field arena and allowed to freely explore for 30 minutes. The animal’s movement was tracked with an overhead video system using EthoVision XT 15 software (Noldus, Leesburg, VA, USA). Total distance travelled, velocity, time spent in the center of the arena, and time spent in the corners of the arena were assessed.

### Statistical Analysis

Initially, sleep architecture data were compared by a three-way repeated-measures (RM) analysis of variance (ANOVA) as follows: total duration of vigilance states (REM, NREM, wake) in 1-hr time bins with treatment (vehicle or PF-04859989) and the time of day (ZT) as within-subject factors and sex (male or female) as a between-subject factor; core body temperature and relative cage activity in 1-hr time bins with treatment (vehicle or PF-04859989) and the time of day (ZT) as within-subject factors and prenatal condition (ECon or EKyn) as a between-subject factor; and NREM and REM onset and transitions between vigilance states with treatment (vehicle or PF-04859989) and phase (light or dark) as within-subject factors and prenatal condition (ECon or EKyn) as a between-subject factor. Significant main effect of treatment and interactions with treatment were analyzed by two-way RM ANOVA as follows: total duration of vigilance states (REM, NREM, wake), number of bouts, and average bout duration in 6-hr time bins with the time of day (ZT) as a within-subject factor; EEG power spectra, NREM delta power and REM theta power, with frequency as a within-subject factor; absolute change in total duration of vigilance states (REM, NREM, wake) and NREM delta spectral power with the time of day (ZT) and frequency as a within-subject factor, respectively; and NREM and REM onset with the prenatal condition (ECon and EKyn) as a within-subject factor. Relative cage activity and core body temperature were analyzed by mixed-effects analysis for each prenatal condition separately. Impact of PF-04859989 treatment on vigilance state duration and relative cage activity was evaluated by calculating the percent change from vehicle. The absolute difference in vigilance state duration and NREM delta spectral power were calculated as a within-subject difference between treatments. Open field data were evaluated by a two-way ANOVA with prenatal condition (ECon or EKyn) and phase (light or dark) as a between-subject factors. In all analyses, where appropriate, significant main effects were followed up with the Fisher’s LSD post hoc analyses. Statistical significance was defined as P < 0.05. All statistical analyses were performed using GraphPad Prism 9.0 software (GraphPad Software, La Jolla, CA, USA). Statistical comparisons and effects sizes are provided in the figure legends.

## Results

To evaluate the impact of reducing KYNA formation on sleep and arousal in adult rats, sleep parameters were assessed in young adult male and female ECon and EKyn offspring after PF-04859989 treatment at ZT 0 or ZT 12. The experimental dose of PF-04859989 was selected based on previously published reports wherein extracellular KYNA levels were substantially reduced across brain regions (25, 41).

### Early Light Phase KAT II Inhibition Enhances Sleep

KYNA levels are elevated during the light phase in EKyn offspring, which may causally contribute to decreased REM duration and number of REM bouts (32, 35, 38). Thus, we presently sought to determine if inhibition of KYNA synthesis via systemic treatment with a KAT II inhibitor would restore REM sleep parameters. When treated with PF-04859989 at ZT 0, REM duration was significantly enhanced across 24 hours in both ECon and EKyn offspring, and analyses separated by sex revealed a significant effect of PF-04859989 in male rats (**Figure 2A, 2B**). Total REM duration was increased by 18-19% during the light phase in ECon, and by 31% during the second half of the light phase in EKyn offspring (**Figure 2C, 2D**), suggesting a rapid enhancement in REM sleep following PF-04859989 treatment.

**Figure 2.**
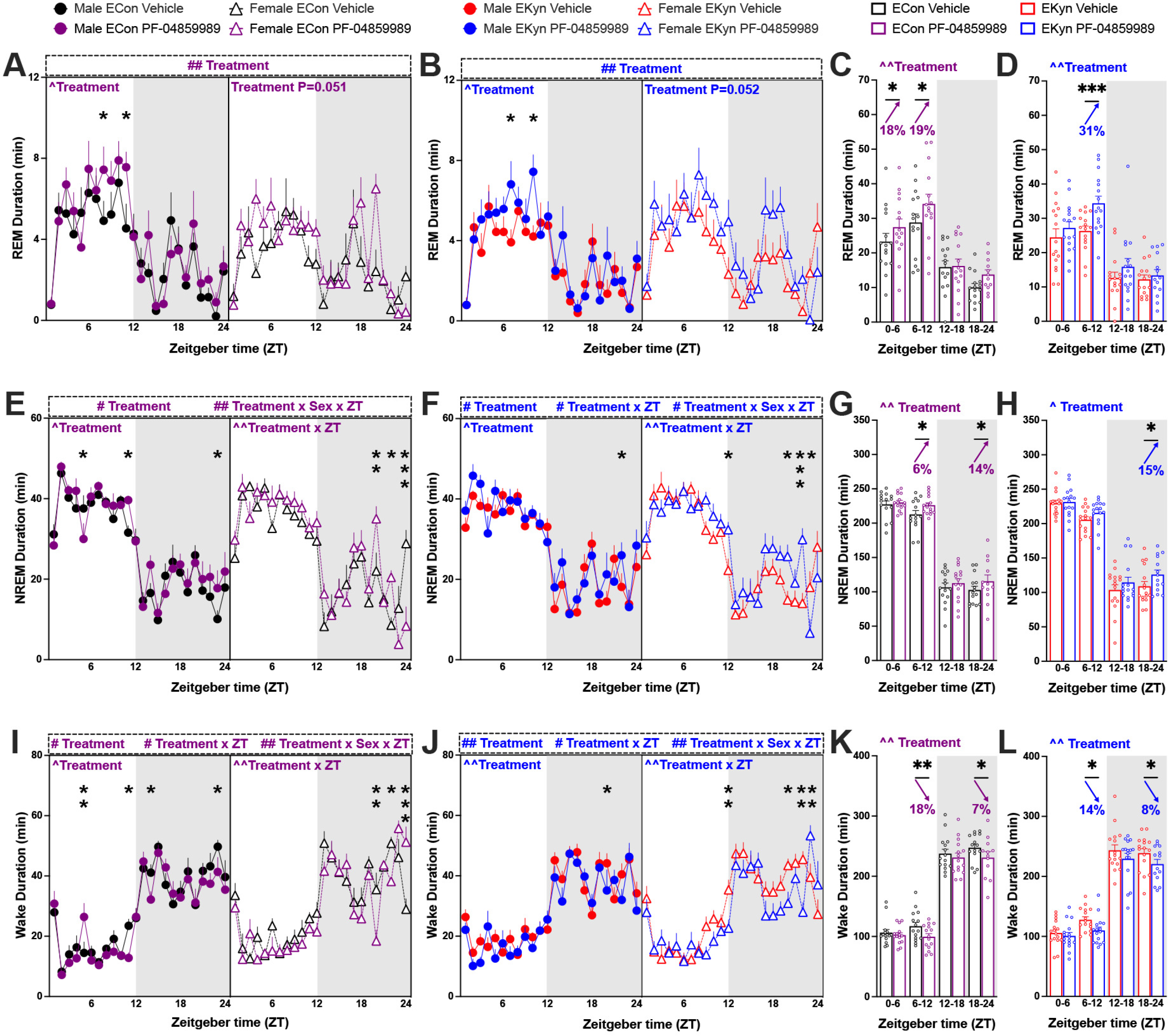
Inhibition of KYNA synthesis at the beginning of light phase promotes REM and NREM sleep and reduces wakefulness in ECon and EKyn offspring. Adult ECon and EKyn offspring were treated with vehicle or PF-04859989 (30 mg/kg) at Zeitgeber time (ZT) 0. **(A)** 1-hr bins of REM duration in male and female ECon (Three-way ANOVA: Treatment effect F_(1, 13)_= 10.00, P<0.01; Two-way ANOVA: Male Treatment effect F_(1, 7)_= 8.938, P<0.05). **(B)** 1-hr bins of REM duration in male and female EKyn (Three-way ANOVA: Treatment effect F_(1, 14)_= 11.76, P<0.01; Two-way ANOVA: Male Treatment effect F_(1, 7)_= 7.211, P<0.05). **(C)** 6-hr bins of REM duration in ECon, sexes combined (Treatment effect F_(1, 14)_= 10.22, P<0.01). **(D)** 6-hr bins of REM duration in EKyn, sexes combined (Treatment effect F_(1, 15)_= 11.15, P<0.01). **(E)** 1-hr bins of NREM duration in male and female ECon (Three-way ANOVA: Treatment effect F_(1, 13)_= 6.135, P<0.05, Treatment x Sex x ZT interaction F_(23, 281)_= 2.005, P<0.01; Two-way ANOVA: Male Treatment effect F_(1, 7)_= 6.976, P<0.05, Female Treatment × ZT interaction F_(23, 120)_= 2.088, P<0.01). **(F)** 1-hr bins of NREM duration in male and female EKyn (Three-way ANOVA: Treatment effect F_(1, 14)_= 8.530, P<0.05, Treatment × ZT interaction F_(23, 310)_= 1.850, P<0.05, Treatment × Sex × ZT interaction F_(23, 310)_= 1.780, P<0.05; Two-way ANOVA: Male Treatment effect F_(1, 7)_= 6.510, P<0.05, Female Treatment × ZT interaction F_(23, 149)_= 2.218, P<0.01). **(G)** 6-hr bins of NREM duration in ECon, sexes combined (Treatment effect F_(1, 14)_= 10.74, P<0.01). **(H)** 6-hr bins of NREM duration in EKyn, sexes combined (Treatment effect F_(1, 15)_= 7.915, P<0.05). **(I)** 1-hr bins of wake duration in male and female ECon (Three-way ANOVA: Treatment effect F_(1, 13)_= 7.459, P<0.05, Treatment × ZT interaction F_(23, 281)_= 1.720, P<0.05, Treatment × Sex × ZT interaction F_(23, 281)_= 2.011, P<0.01; Two-way ANOVA: Male Treatment effect F_(1, 7)_= 6.445, P<0.05, Female Treatment × ZT interaction F_(23, 120)_= 2.114, P<0.01). **(J)** 1-hr bins of wake duration in male and female EKyn (Three-way ANOVA: Treatment effect F_(1, 14)_= 12.07, P<0.01, Treatment × ZT interaction F_(23, 310)_= 1.755, P<0.05, Treatment × Sex × ZT interaction F_(23, 310)_= 1.875, P<0.01; Two-way ANOVA: Male Treatment effect F_(1, 7)_= 12.30, P<0.01, Female Treatment × ZT interaction F_(23, 149)_= 2.235, P<0.01). **(K)** 6-hr bins of wake duration in ECon, sexes combined (Treatment effect F_(1, 14)_= 11.98, P<0.01). **(L)** 6-hr bins of wake duration in EKyn, sexes combined (Treatment effect F_(1, 15)_= 11.05, P<0.01). Data are mean ± SEM. Percent change from vehicle treatment calculations are shown by arrows. Three-way RM ANOVA: # P<0.05, ## P<0.01. Two-way RM ANOVA: ^P<0.05, ^^P<0.01 with Fisher’s LSD post hoc test: *P<0.05, **P<0.01, ***P<0.001. N = 12-16 per group.

Over 24 hours, PF-04859989 treatment significantly impacted NREM duration in ECon and EKyn offspring (**Figure 2E, 2F**). Total NREM duration was increased during the latter part of both light and dark phases in ECon, +6% and +14%, respectively, and +15% during the second half of the dark phase in EKyn offspring (**Figure 2G, 2H**). Sleep architecture is characterized by the number and length of bouts of REM and NREM sleep. PF-04859989 treatment significantly impacted NREM bout number in ECon and EKyn rats, wherein NREM bout number increased by 17% during the late dark phase (**Supplementary Figure 1**).

Each vigilance state comprises bouts of various lengths and bouts sustained for a longer time indicate a firmly consolidated vigilance state. We noted a significant treatment x ZT interaction for average duration of both REM and NREM bouts with a phase-specific effects of PF-04859989 treatment. Average REM bout duration in EKyn rats was increased by 34% during the first half and decreased by 28% during the second half of the light phase, while average NREM bout duration was increased by 13% during early light phase in ECon rats (**Supplementary Figure 1**).

As REM and NREM sleep were enhanced, wakefulness was reduced. With PF-04859989 treatment at the beginning of the light phase subjects were awake less time across 24 hours, determined by a main effect of treatment, a treatment x ZT interaction, as well as an interaction between treatment x sex x ZT (**Figure 2I, 2J**). Male and female ECon and EKyn offspring exhibited reduced wake duration during the second half of light phase, −15% and −14% respectively, and dark phase, −7% and −8%, respectively (**Figure 2K, 2L**). Taken together, treatment of adult offspring with PF-04859989 at ZT 0 induced enhanced somnolence.

### KAT II Inhibition Enhances NREM Delta Spectral Power

Delta power during NREM sleep correlates strongly with sleep quality (42), and theta oscillations during REM sleep are implicated in memory consolidation (43, 44). Therefore, we next sought to detect if acutely inhibiting KAT II would be sufficient to drive changes in EEG waves within low-frequency oscillation bands, delta (0-4 Hz) and theta (4-8 Hz). When PF-04859989 was administered at the beginning of the light phase, we found significant effects of KAT II inhibition on delta spectral power during both the light and dark phases, but with subtle sex-dependent differences. In male ECon offspring, a treatment x frequency interaction pointed to a significant increase in NREM delta power immediately after the treatment during the light phase (**Figure 3A**), and the effect of PF-04859989 treatment lasted through the dark phase (**Figure 3B**). In adult male EKyn rats, PF-04895589 treatment significantly elevated the low-frequency delta bands only during the light phase (**Figure 3C, 3D**). Among female offspring, NREM power in the lower frequency range was enhanced, as determined by a treatment x frequency interaction in ECon rats during the light phase alone (**Figure 3E, 3F, 3G, 3H**). Analyses of theta spectral power after acute KYNA reduction at ZT 0 showed an interaction between treatment x frequency and alterations within REM theta power in EKyn male rats during the light phase (**Supplementary Figure 2**).

**Figure 3.**
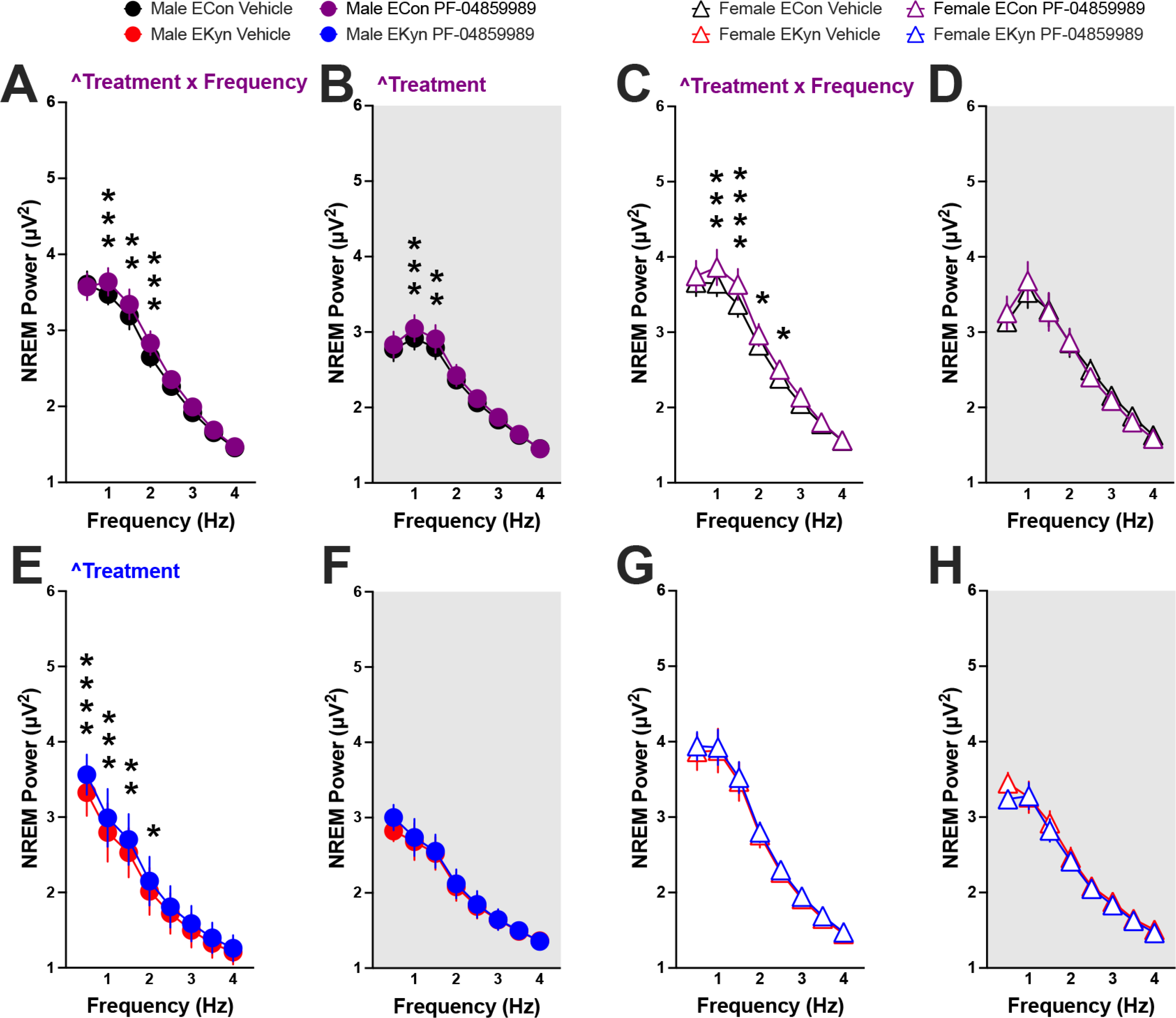
Reduction in KYNA levels at the beginning of light phase enhances NREM delta spectral power in ECon and EKyn offspring. Adult ECon and EKyn offspring were treated with vehicle or PF-04859989 (30 mg/kg) at Zeitgeber time (ZT) 0. **(A)** Male ECon during light phase (Treatment x Frequency interaction F_(7, 49)_= 2.892, P<0.05). **(B)** Male ECon during dark phase (Treatment effect F_(1, 7)_= 6.678, P<0.05). **(C)** Female ECon during light phase (Treatment x Frequency interaction F_(7, 42)_= 2.897, P<0.05). **(D)** Female ECon during dark phase. **(E)** Male EKyn during light phase (Treatment effect F_(1, 7)_= 9.155, P<0.05). **(F)** Male EKyn during dark phase. **(G)** Female EKyn during light phase. **(H)** Female EKyn during dark phase. Data are mean ± SEM. Two-way RM ANOVA: ^P<0.05 with Fisher’s LSD post hoc test: *P<0.05, **P<0.01, ***P<0.001, ****P<0.0001. N = 7-8 per group.

### Dark Phase KAT II Inhibition Promotes Sleep During the Subsequent Light Phase

Administration of KAT II inhibitor at ZT 12 elicited a treatment x ZT interaction across all three vigilance states in ECon offspring, with total REM and NREM duration increased during the first half of the successive light phase by 19% and 14%, respectively (**Figure 4A, 4B**). Increase in sleep duration between ZT 0 and ZT 6 during the succeeding light phase coincided with 28% reduction in wake duration (**Figure 4C**). During the latter part of the subsequent light phase, between ZT 6 and ZT 12, NREM duration was reduced by 11%, and wake duration increased by 21% in ECon subjects (**Figure 4B, 4C**). Assessment of sleep architecture detected that the number of wake bouts increased with PF-04859989 treatment in ECon rats between ZT 0 and ZT 6, despite spending less time awake during this period (**Supplementary Figure 3**). No changes in NREM power spectra were found with PF-04859989 treatment at ZT 12 (**Figure 4D, 4E**).

**Figure 4.**
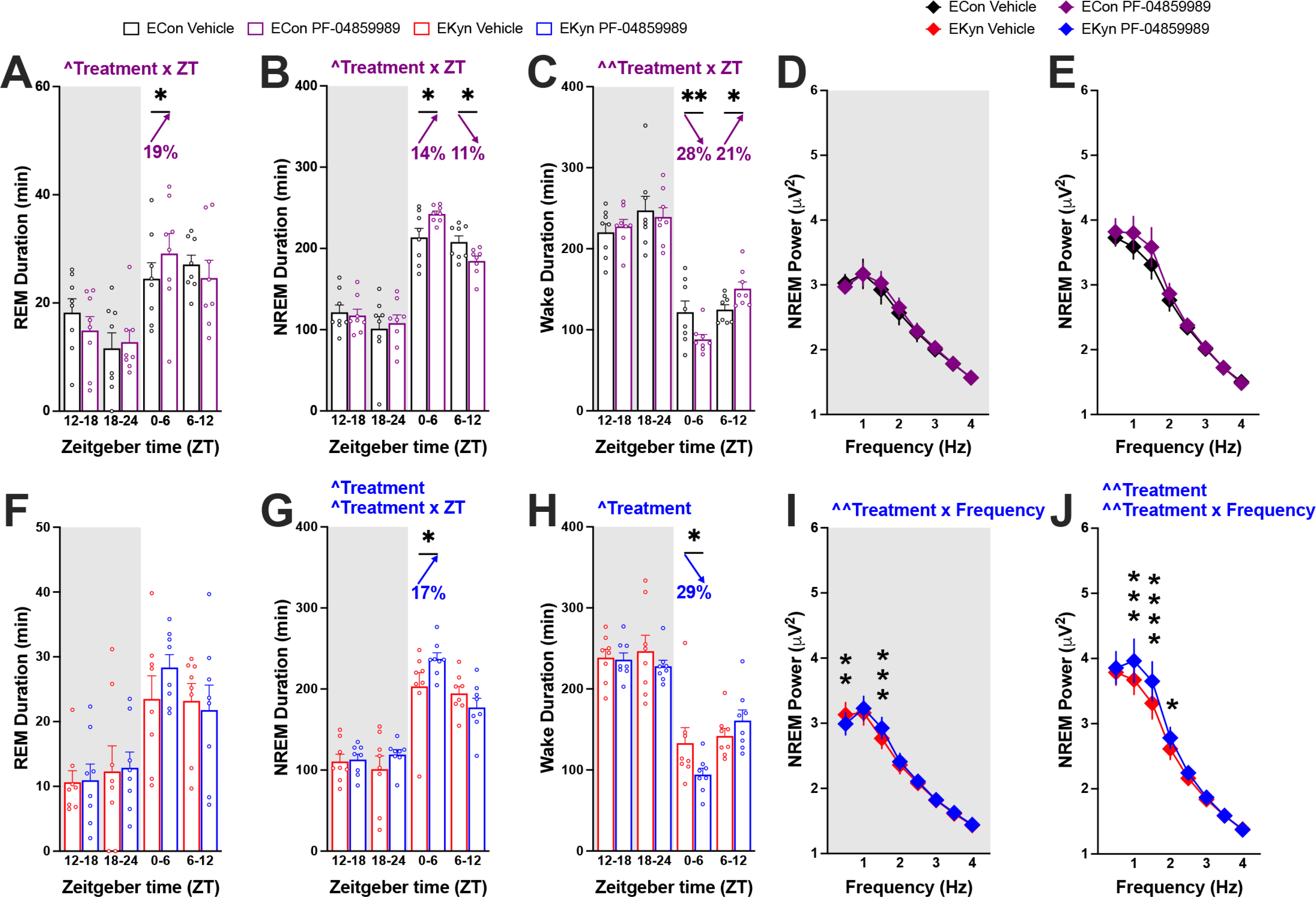
KAT II inhibition at the beginning of dark phase improves sleep during the subsequent light phase in ECon and EKyn offspring. Adult ECon and EKyn offspring were treated with vehicle or PF-04859989 (30 mg/kg) at Zeitgeber time (ZT) 12. **(A)** 6-hr bins of REM duration in ECon, sexes combined (Treatment x ZT interaction F_(3, 21)_= 3.451, P<0.05). **(B)** 6-hr bins of NREM duration in ECon, sexes combined (Treatment x ZT interaction F_(3, 21)_= 4.355, P<0.05). **(C)** 6-hr bins of wake duration in ECon, sexes combined (Treatment x ZT interaction F_(3, 21)_= 5.087, P<0.01). **(D)** NREM delta spectral power during dark phase in ECon, sexes combined. **(E)** NREM delta spectral power during light phase in ECon, sexes combined. **(F)** 6-hr bins of REM duration in EKyn, sexes combined. **(G)** 6-hr bins of NREM duration in EKyn, sexes combined (Treatment effect F_(1, 7)_= 5.594, P=0.05, Treatment x ZT interaction F_(3, 21)_= 3.290, P<0.05). **(H)** 6-hr bins of wake duration in EKyn, sexes combined (Treatment effect F_(1, 7)_= 6.926, P<0.05). **(I)** NREM delta spectral power during dark phase in EKyn, sexes combined (Treatment x Frequency interaction F_(7, 49)_= 3.861, P<0.01). **(J)** NREM delta spectral power during light phase in EKyn, sexes combined (Treatment effect F_(1, 7)_= 13.29, P<0.01, Treatment x Frequency interaction F_(7, 49)_= 3.050, P<0.01). Data are mean ± SEM. Percent change from vehicle treatment calculations are shown by arrows. Two-way RM ANOVA: ^P<0.05, ^^P<0.01 with Fisher’s LSD post hoc test: *P<0.05, **P<0.01, ***P<0.001, ****P<0.0001. N = 8 per group.

Treatment of EKyn offspring at the start of the dark phase with PF-04859989 produced similar impacts as in ECon. PF-04859989 treatment at ZT 12 in EKyn rats elevated NREM duration by 17% and decreased wake duration by 29% during the first half of the consecutive light phase without altering total REM duration (**Figure 4F, 4G, 4H**). Sleep architecture, namely the number of NREM and wake bouts, was significantly impacted with PF-04859989 treatment (**Supplementary Figure 3**). In contrast to dark phase PF-04859989 treatment in ECon offspring, NREM delta power was significantly impacted when EKyn offspring were treated with PF-04859989 at ZT 12, noted as a statistically significant treatment x frequency interaction throughout both dark and light phases (**Figure 4I, 4J**).

### Administration of KAT II Inhibitor Impacts Sleep-Wake Transitions

Changes in sleep architecture were further assessed by evaluating transitions between vigilance states. PF-04859989 treatment increased the number of REM to NREM transitions when the drug was administered at ZT 0. With treatment at ZT 12, the number of transitions from wake to NREM sleep and conversely from NREM sleep to wake were significantly increased with PF-04859989 (**Table 1**). The elevated number of transitions from NREM to wake supports our findings that the number of wake bouts is increased by PF-04859989 treatment without a change in the number of NREM bouts (**Supplementary Figure 3**). Sleep onset latency, evaluating sleep propensity (45), was unchanged within the immediate light or dark phase after PF-04859989 treatment, however REM onset was during the subsequent dark phase with ZT 0 treatment (**Table 1**).

**Table 1.**
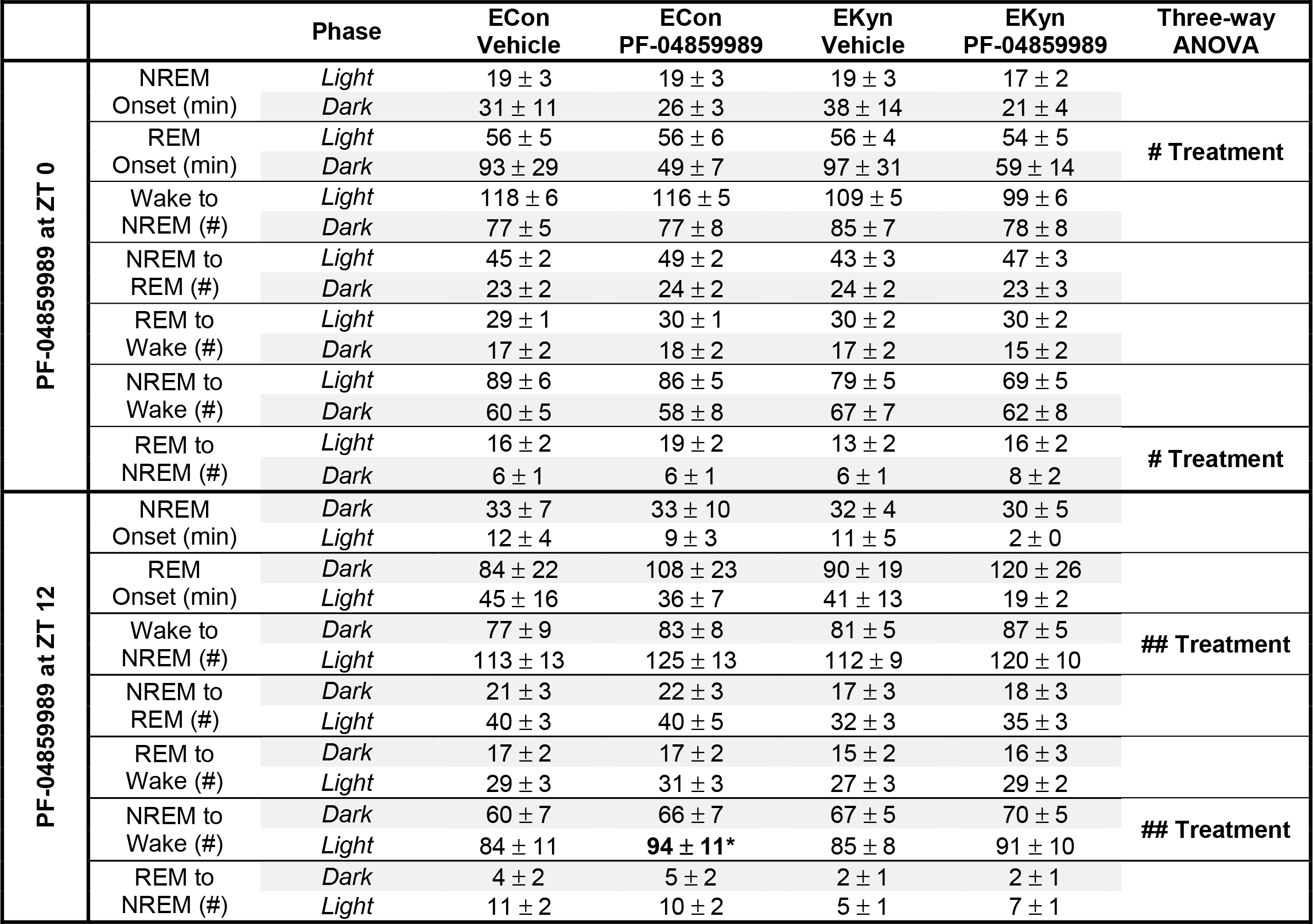
KAT II inhibitor, PF-04859989, alters sleep onset and transitions between vigilance states. Data are mean ± SEM. Three-way RM ANOVA: # P<0.05, ## P<0.01. Two-way RM ANOVA with Fisher’s LSD post hoc test: *P<0.05. N = 8-16 per group.

### Sex-Dependent Novelty-induced Exploration

EKyn offspring pose sex-dependent patterns in behavior including reduced home cage activity (38). To further characterize these phenotypes, we evaluated basal locomotion and exploration in the open field during both light and dark phases in both sexes of offspring. Exposure to the open field arena induced novelty-induced exploration in a phase-dependent manner. Regardless of prenatal treatment, females had increased ambulation (**Figure 5A**), and higher average velocity (**Figure 5B**) in comparison to males across 24 hours. Of note, exploration of the center of the arena was significantly greater during dark phase compared to light phase testing in males (**Figure 5C**). Similarly, the percent of time spent in the corners of the arena was significantly lower during dark phase compared to light phase testing, in both sexes and prenatal conditions (**Figure 5D**). We did not determine any EKyn-dependent deficits in open field exploration across measures, however female EKyn spent more time exploring the center of the testing arena compared to EKyn males during the light phase.

**Figure 5.**
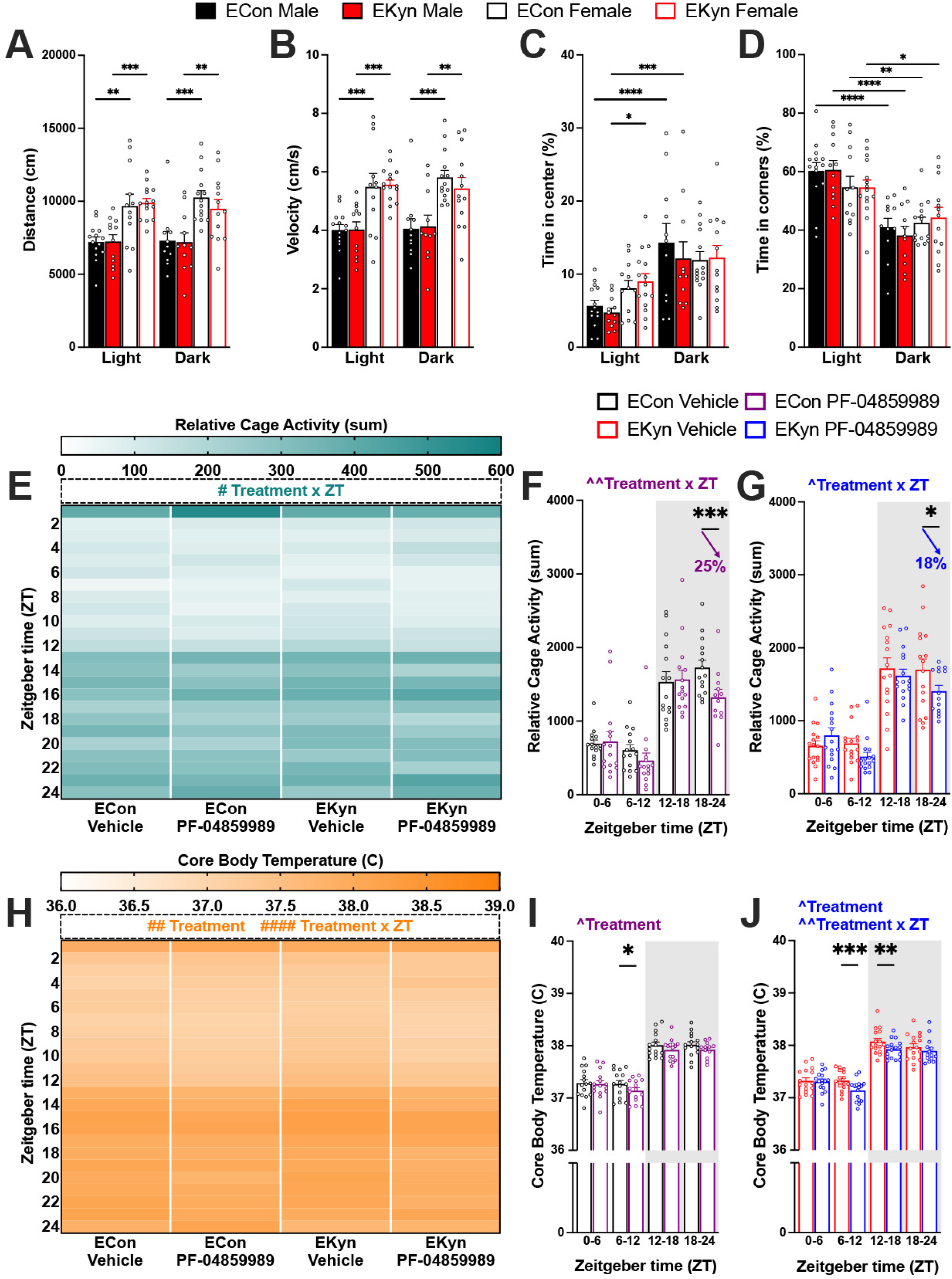
Administration of KAT II inhibitor at beginning of light phase reduces activity and temperature in ECon and EKyn offspring who exhibit sex differences in locomotor activity. Male and female offspring were tested in the open field test during the light or dark phase: **(A)** Distance travelled, **(B)** Velocity, **(C)** Percent of time spent in center, **(D)** Percent of time spent in corners. Adult ECon and EKyn offspring were treated with vehicle or PF-04859989 (30 mg/kg) at Zeitgeber time (ZT) 0. **(E)** 1-hr bins of relative cage activity, sexes combined (Three-way ANOVA: Treatment x ZT interaction F_(23, 487)_= 1.825, P<0.05). **(F)** 6-hr bins of relative cage activity in ECon, sexes combined (Treatment x ZT interaction F_(3, 39)_= 5.621, P<0.01). **(G)** 6-hr bins of relative cage activity in EKyn, sexes combined (Treatment x ZT interaction F_(3, 43)_= 3.175, P<0.05). **(H)** 1-hr bins of core body temperature, sexes combined (Three-way ANOVA: Treatment effect F_(1, 29)_= 9.403, P<0.01, Treatment x ZT interaction F_(23, 480)_= 2.978, P<0.0001). **(I)** 6-hr bins of core body temperature in ECon, sexes combined (Treatment effect F_(1, 14)_= 7.810, P<0.05). **(J)** 6-hr bins of core body temperature in EKyn, sexes combined (Treatment effect F_(1, 15)_= 7.507, P<0.05, Treatment x ZT interaction F_(3, 43)_= 6.280, P<0.01). Data are mean ± SEM. Percent change from vehicle treatment calculations are shown by arrows. Three-way ANOVA: # P<0.05, ## P<0.01, #### P<0.0001. Two-way RM ANOVA: ^P<0.05, ^^P<0.01 with Fisher’s LSD post hoc test: *P<0.05, **P<0.01, ***P<0.001, ****P<0.0001. N = 11-16 per group.

### Reduced Cage Activity and Body Temperature with PF-04859989 Treatment

Active wakefulness, characterized by limb movements and locomotor activity, differs from quiescent wake behavior (46). PF-04859989 administered at the beginning of the light phase significantly impacted cage activity, noted as a treatment x ZT interaction (**Figure 5E**). Relative cage activity was decreased after PF-04859989 in both prenatal groups during the second half of the dark phase (**Figure 5F, 5G**). Rats typically exhibit elevated core temperature and increased motor activity during the dark phase (47). We found that PF-04859989 also impacted core body temperature, noted as an interaction between treatment and ZT (**Figure 5H**). In ECon rats, body temperature decreased only in the latter half of the light phase (**Figure 5I**), whereas in EKyn offspring the decrease in core body temperature was sustained between ZT 6 and ZT 18 (**Figure 5J**). Treatment with PF-04859989 at the beginning of the dark phase produced no significant differences in relative cage activity or core body temperature (**Supplementary Figure 4**).

### Enhanced improvements in sleep architecture with PF-04859989 treatment at the start of the light phase

To compare the effectiveness of PF-04859989 administration between the start of the light or dark phase, we evaluated the absolute changes in duration of the three vigilance states. Notably, during the first 12 hours post-drug administration, we noted stark differences in the enhancement of REM duration (**Supplementary Figure 6A**). Changes in NREM duration and wake duration, induced by PF-04859989 treatment, were also impacted by the time of treatment, but major distinctions were found 12 hours after drug administration (**Supplementary Figure 6B, 6C**). Absolute changes in NREM delta spectral power were also evaluated, comparing the first 12 hours after PF-04859989 treatment and the subsequent 12 hours. We observed consistent improvement in NREM power spectrum within the first 12 hours regardless of PF-04859989 administration at ZT 0 or ZT 12 (**Supplementary Figure 6D**). Spectral power, in the lowest delta frequencies between 0.5–2 Hz, was notably differentially impacted between 12 and 24 hours after PF-04859989 administration (**Supplementary Figure 6E**).

## Discussion

Our present study is the first to signify that acute KAT II inhibition improves sleep physiology and architecture, thereby suggesting that reducing brain KYNA alleviates sleep disturbances. KAT II inhibitor PF-04859989 effectively enhanced sleep parameters in both ECon and EKyn offspring of both sexes. Our findings contribute to the recent growing body of literature showing that the pivotal enzymes within the KP serve as effective physiological treatment targets (10, 48).

Tryptophan metabolism has long been implicated in modulating sleep, namely through its formation of the circadian hormone melatonin (49, 50). Yet, a role for KP metabolites, specifically KYNA, in regulating sleep-wake behavior has only recently been discovered. Acutely elevating de novo production of brain KYNA showed prompt reduction in REM sleep, including alterations in REM architecture (39). Similarly, REM sleep dynamics are disrupted in young adult EKyn offspring, wherein elevated brain KYNA levels are postulated to disrupt sleep and arousal patterns compared to counterpart controls. EKyn males have reduced duration of REM sleep concomitant with failing to initiate REM episodes, while the number of both NREM and wake bouts was lower in EKyn females compared to controls (38). We presently leverage the strengths of our EKyn preclinical model to test the novel hypothesis that reducing brain KYNA can impact sleep duration and quality, and aim to place further attention on the KP as a targetable metabolic pathway to improve sleep homeostasis.

KYNA is an endogenous antagonist of the glutamatergic N-methyl-D-aspartate (NMDA) receptors and alpha-7 nicotinic acetylcholine receptors (α7nAChR)(16, 51). Cholinergic transmission is mediated by activation of the presynaptic α7nAChRs on glutamatergic neurons and the postsynaptic α7nAChRs on parvalbumin GABA (gamma-aminobutyric acid)-ergic interneurons (52, 53). Decrease in KYNA levels is postulated to physiologically enhance cholinergic signaling across the brain and stimulate release of glutamate (15). As the generation of REM sleep and sleep stability is promoted by excitatory glutamatergic neurotransmission (46, 54), we can presently speculate that global reduction in KYNA contributes to the stability of REM sleep. In the EKyn rats, extracellular glutamate levels are reduced during the light phase (35), and KAT II inhibition effectively restores glutamate to control levels (37). Further exploring circuit level modulation from the brainstem pontine areas to the cerebral cortex and the hippocampus to regulate REM sleep stability is warranted in future studies (55–57).

Demand for novel, efficacious sleep therapeutics has increased substantially in the recent years. Treatments that improve sleep onset latency, maintenance, efficiency, slow wave power, and overall sleep quality could ameliorate commonly reported sleep problems in SCZ and BPD patients (58–67). Unfortunately, many commonly prescribed antipsychotic drugs, along with adjunct antidepressant therapeutic approaches, suppress sleep further (68, 69). Our present findings are translationally noteworthy as KAT II inhibition effectively improves sleep outcomes in both EKyn, our neurodevelopmental insult paradigm, and the counterpart controls, ECon. Single administration of KAT II inhibitor was sufficient to elicit recovered sleep architecture in EKyn rats to the levels that are observed in ECon rats for the subsequent 24-hour period. Evaluation of the time of day of PF-04859989 treatment concluded that treatment at the beginning of the light phase triggers immediate improvements in sleep behavior, while treatment at the beginning of the dark phase deferred outcomes until the subsequent light phase. Notably, the ability of PF-04859989 to maintain entrainment of the light-dark cycle in adult rodents further supports the translational potential of pharmacological approaches targeting KAT II.

In addition to promoting sleep duration, the KAT II inhibitor enhanced NREM delta power, potentially advancing overall sleep quality. Delta spectral power indicates homeostatic sleep drive, but it is regulated independently of sleep duration (70). Delta waves of NREM sleep occur when neurons originating from extensive areas of the cortex repeatedly transit between a hyperpolarized and depolarized state (71, 72). These slow cortical oscillations couple and synchronize with sharp-wave ripples generated in the hippocampus to facilitate memory consolidation and various cognitive functions (73–75). PF-04859989 treatment enhanced NREM spectral power dynamically in a sex- and phase-dependent manner presently. This finding is informative for the development of targeted therapeutic strategies to enhance sleep quality, and in tandem have the potential to enhance cognitive function (25).

Cognitive deficits and memory impairments are lingering symptoms that persist in individuals with severe psychiatric illnesses albeit antipsychotic therapy (76–79). Clinical studies also point to sex differences in cognitive performance in individuals with SCZ. Notably across several studies, cognitive deficits in males are more adverse and their response to various antipsychotic medications is less effective (80–85). Our EKyn model is highly translationally relevant, as we have determined sex differences in cognitive outcomes that align with the aforementioned clinical findings. Male EKyn offspring are more adversely impacted in learning and cognitive flexibility in behavioral tasks compared to female offspring (32). KAT II inhibitor PF-04859989 promotes sleep quality, irrespective of sex, and thus we speculate that PF-04859989 treatment of EKyn offspring may also effectively improve cognitive performance, as in behavioral experiments with KAT II inhibitor BFF-816 in EKyn offspring (37).

Sleep quality may also contribute to avolition, considered a core negative symptom in individuals with SCZ and BPD (86–89). To evaluate if sleep disturbances contribute to a lack of motivation, slow movement, and fatigue, exploratory and locomotor behaviors were evaluated, yet we determined no differences between ECon and EKyn offspring. We did, however, note higher mobility in female rats, as reported by others (90). Female behavioral and physiological processes, including cognition and sleep, are greatly impacted by the hormonal status and the estrous cycle (91, 92) and the presentation of symptoms in females with SCZ is also often at a later age than males (93). To comprehensively define a role for KYNA in the timeline of sleep and affective symptoms, future preclinical and clinical studies will be imperative (89, 94, 95).

In closing, we presently place critical attention on a novel targeted therapeutic strategy for overcoming sleep disorders. Acute KAT II inhibition to reduce KYNA is shown to enhance REM and NREM sleep, and restore sleep stability in EKyn, our neurodevelopmental insult model. Improvement in sleep stability places much needed attention on a targeted approach which may lead to the development of potential therapeutic interventions in individuals combatting sleep disruptions alongside neuropsychiatric and neurodevelopmental disorders.

## Acknowledgements

The authors would like to thank Audrey Ditty, Katherine Rentschler, and Nathan Wagner for their technical contributions to this work. This research was supported by National Institutes of Health Grant Nos. NIH R01 NS102209 (AP), P50 MH103222 (AP), and P20 RR-016461 (HV).

