## Supplemental Materials for "Kynurenine aminotransferase II inhibition promotes sleep and rescues impairments induced by neurodevelopmental insult"

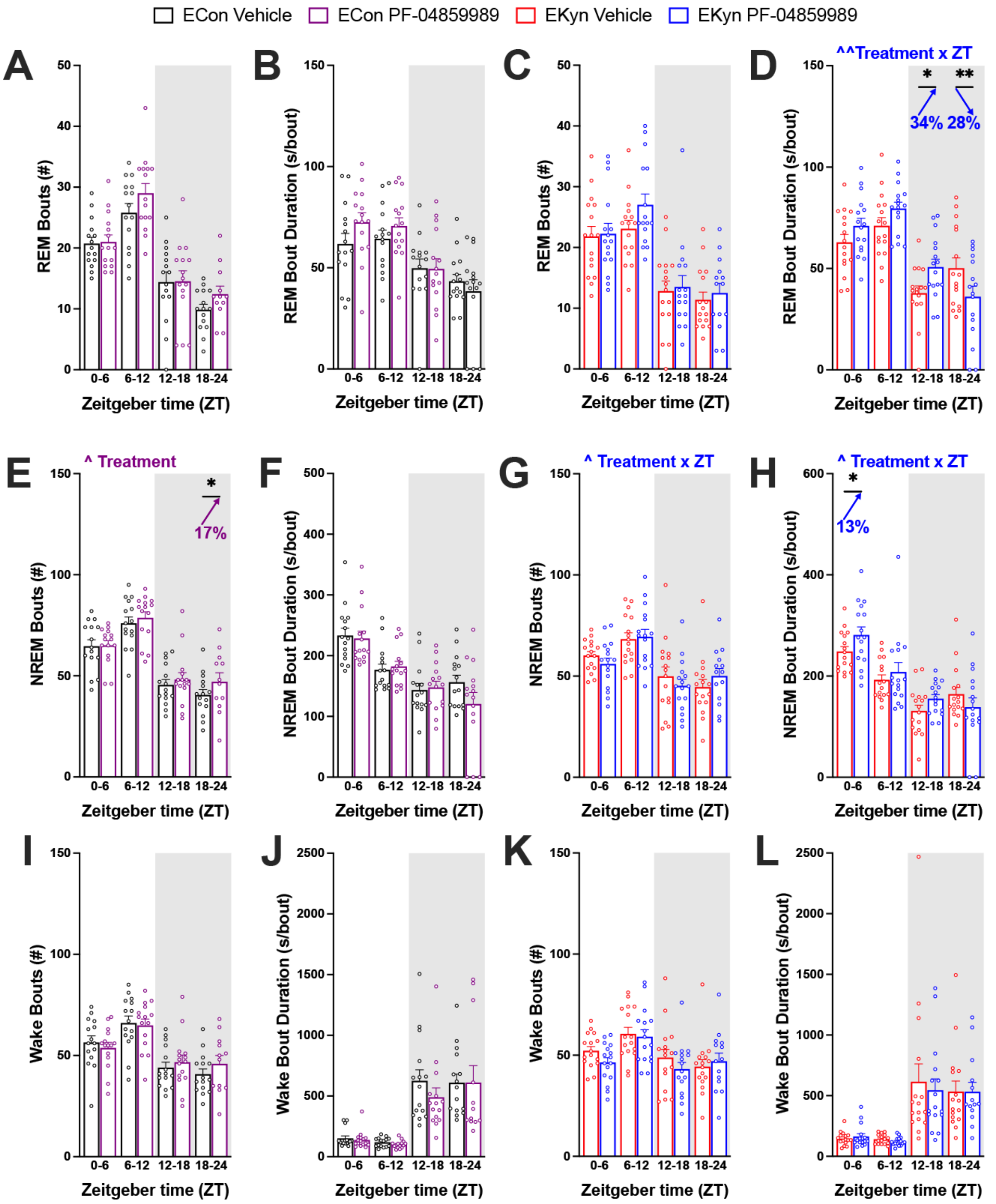


**Supplementary Figure 1. Sleep architecture following inhibition of KYNA synthesis at the beginning of light phase.** Adult ECon and EKyn offspring were treated with vehicle or PF-04859989 (30 mg/kg) at Zeitgeber time (ZT) 0. **(A)** 6-hr bins of number of REM bouts in ECon, sexes combined. **(B)** 6-hr bins of average REM bout duration in ECon, sexes combined. **(C)** 6-hr bins of number of REM bouts in EKyn, sexes combined. **(D)** 6-hr bins of average REM bout duration in EKyn, sexes combined (Treatment x ZT interaction $F_{(3, 45)}$= 6.024, P<0.01). **(E)** 6-hr bins of number of NREM bouts in ECon, sexes combined. **(F)** 6-hr bins of average NREM bout duration in ECon, sexes combined. **(G)** 6-hr bins of number of NREM bouts in EKyn, sexes combined (Treatment x ZT interaction $F_{(3, 43)}$= 3.471, P<0.05). **(H)** 6-hr bins of average NREM bout duration in EKyn, sexes combined (Treatment x ZT interaction $F_{(3, 45)}$= 3.973, P<0.05). **(I)** 6-hr bins of number of wake bouts in ECon, sexes combined. **(J)** 6-hr bins of average wake bout duration in ECon, sexes combined. **(K)** 6-hr bins of number of wake bouts in EKyn, sexes combined. **(L)** 6-hr bins of average wake bout duration in EKyn, sexes combined. Data are mean ± SEM. Percent change from vehicle treatment calculations are shown by arrows. Two-way RM ANOVA: ^P<0.05, ^^P<0.01 with Fisher's LSD post hoc test: *P<0.05, **P<0.01. N = 12-16 per group.

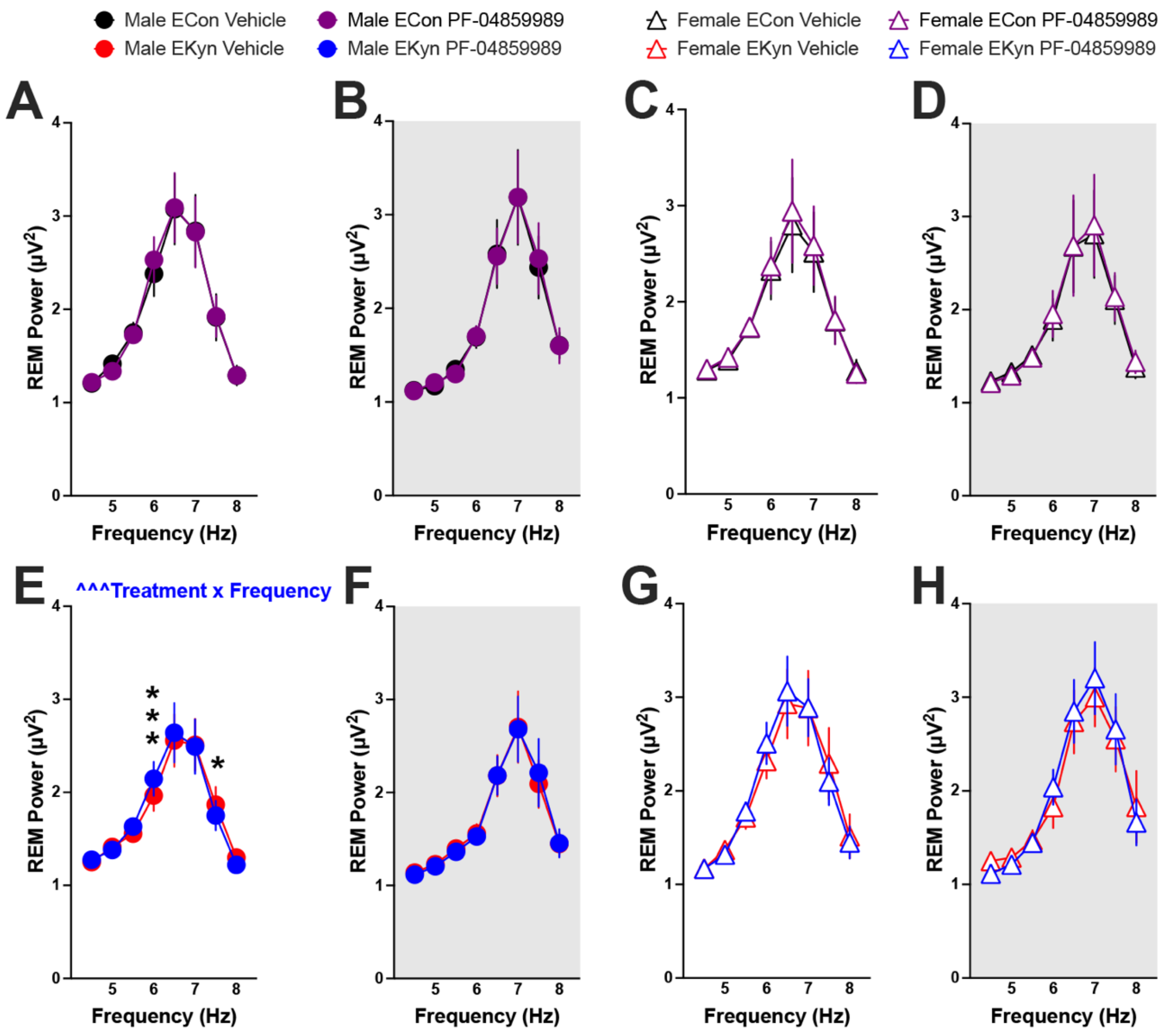


**Supplementary Figure 2. REM theta spectral power after reduction in KYNA levels at the beginning of light phase.** Adult ECon and EKyn offspring were treated with vehicle or PF-04859989 (30 mg/kg) at Zeitgeber time (ZT) 0. **(A)** Male ECon during light phase. **(B)** Male ECon during dark phase. **(C)** Female ECon during light phase. **(D)** Female ECon during dark phase. **(E)** Male EKyn during light phase (Treatment x Frequency interaction $F_{(7, 42)}$= 4.738, P<0.001). **(F)** Male EKyn during dark phase. **(G)** Female EKyn during light phase. **(H)** Female EKyn during dark phase. Data are mean ± SEM. Two-way RM ANOVA: ^^^P<0.001 with Fisher's LSD post hoc test: *P<0.05, ***P<0.001. N = 7-8 per group.

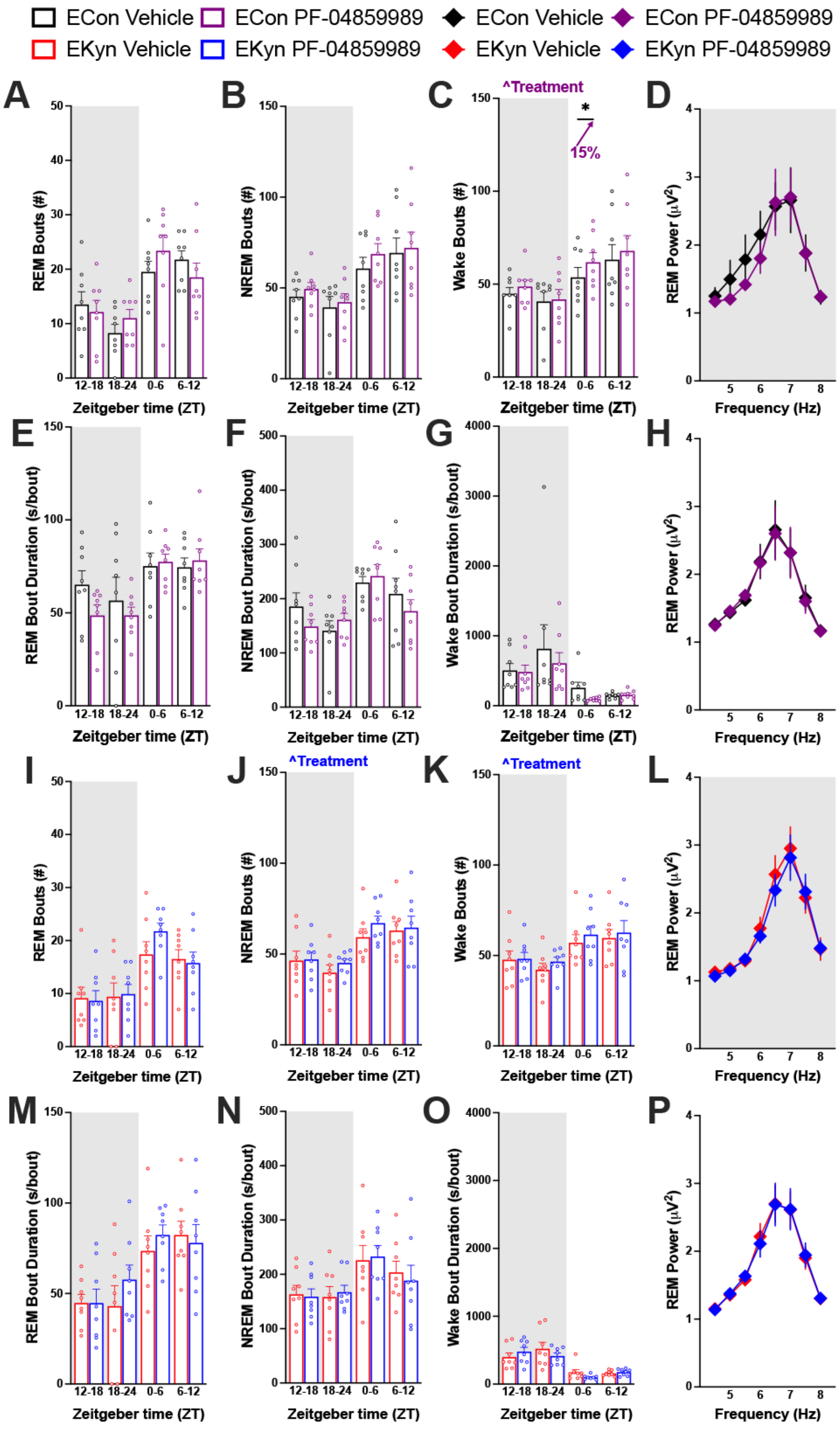

ECon Vehicle
ECon PF-04859989
EKyn Vehicle
EKyn PF-04859989
A
B
C
D
E
F
G
H
I
J
K
L
M
N
O
P
^Treatment
*
15%
REM Bouts (#)
NREM Bouts (#)
Wake Bouts (#)
REM Power (μV²)
REM Bout Duration (s/bout)
NREM Bout Duration (s/bout)
Wake Bout Duration (s/bout)
Zeitgeber time (ZT)
Frequency (Hz)
12-18
18-24
0-6
6-12

**Supplementary Figure 3. Sleep architecture following KAT II inhibition at the beginning of dark phase.** Adult ECon and EKyn offspring were treated with vehicle or PF-04859989 (30 mg/kg) at Zeitgeber time (ZT) 12. **(A)** 6-hr bins of number of REM bouts in ECon, sexes combined. **(B)** 6-hr bins of number of NREM bouts in ECon, sexes combined. **(C)** 6-hr bins of number of wake bouts in ECon, sexes combined (Treatment effect $F_{(1, 7)}$= 6.334, P<0.05). **(D)** REM theta spectral power during dark phase in ECon, sexes combined. **(E)** 6-hr bins of average REM bout duration in ECon, sexes combined. **(F)** 6-hr bins of average NREM bout duration in ECon, sexes combined. **(G)** 6-hr bins of average wake bout duration in ECon, sexes combined. **(H)** REM theta spectral power during light phase in ECon, sexes combined. **(I)** 6-hr bins of number of REM bouts in EKyn, sexes combined. **(J)** 6-hr bins of number of NREM bouts in EKyn, sexes combined (Treatment effect $F_{(1, 7)}$= 6.975, P<0.05). **(K)** 6-hr bins of number of wake bouts in EKyn, sexes combined (Treatment effect $F_{(1, 7)}$= 5.826, P<0.05). **(L)** REM theta spectral power during dark phase in EKyn, sexes combined. **(M)** 6-hr bins of average REM bout duration in EKyn, sexes combined. **(N)** 6-hr bins of average NREM bout duration in EKyn, sexes combined. **(O)** 6-hr bins of average wake bout duration in EKyn, sexes combined. **(P)** REM theta spectral power during light phase in EKyn, sexes combined. Data are mean ± SEM. Percent change from vehicle treatment calculations are shown by arrows. Two-way RM ANOVA: ^P<0.05 with Fisher's LSD post hoc test: *P<0.05. N = 8 per group.

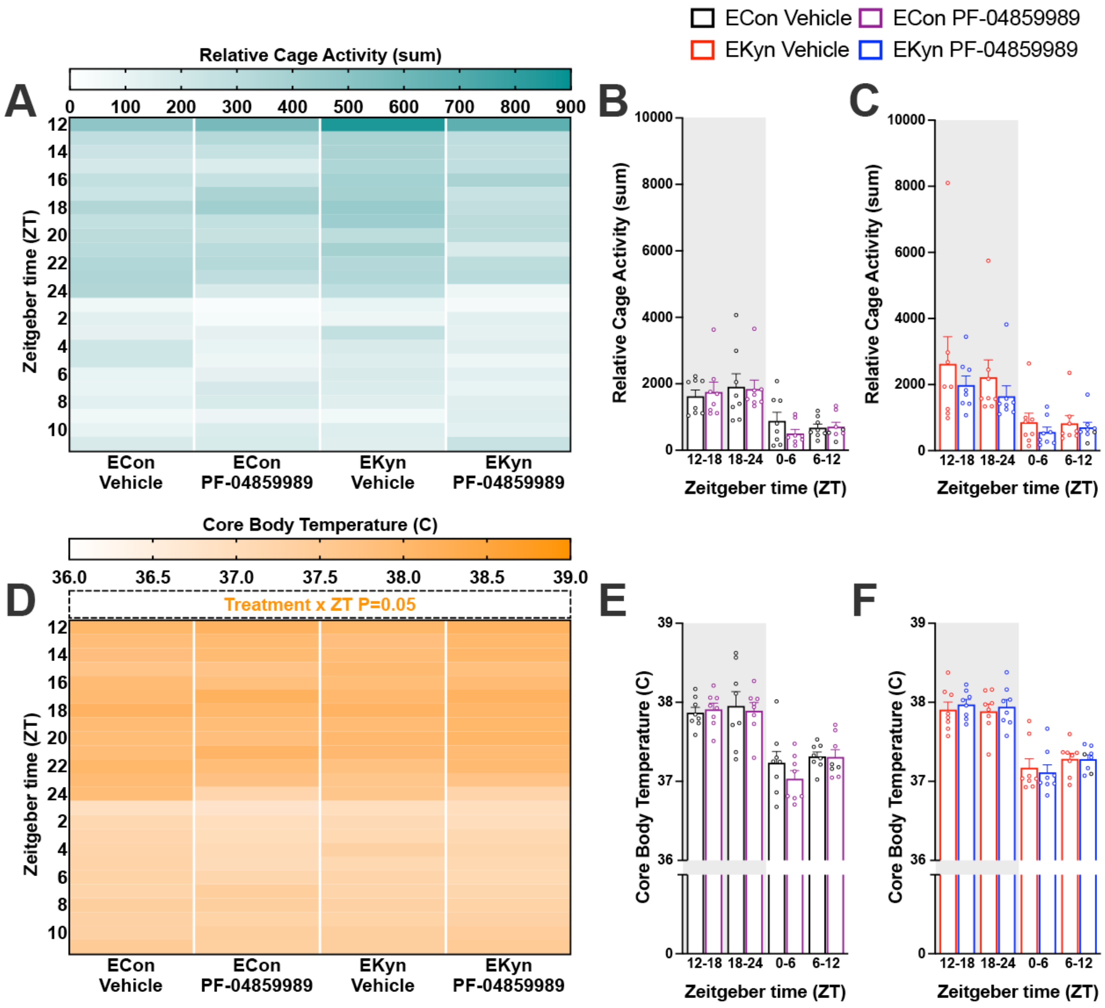


**Supplementary Figure 4. Activity and temperature after administration of KAT II inhibitor at the beginning of dark phase.** Adult ECon and EKyn offspring were treated with vehicle or PF-04859989 (30 mg/kg) at Zeitgeber time (ZT) 12. **(A)** 1-hr bins of relative cage activity, sexes combined. **(B)** 6-hr bins of relative cage activity in ECon, sexes combined. **(C)** Relative cage activity in EKyn, sexes combined. **(D)** 1-hr bins of core body temperature, sexes combined. **(E)** 6-hr bins of core body temperature in ECon, sexes combined. **(F)** Core body temperature in EKyn, sexes combined. Data are mean ± SEM. N = 11-16 per group.

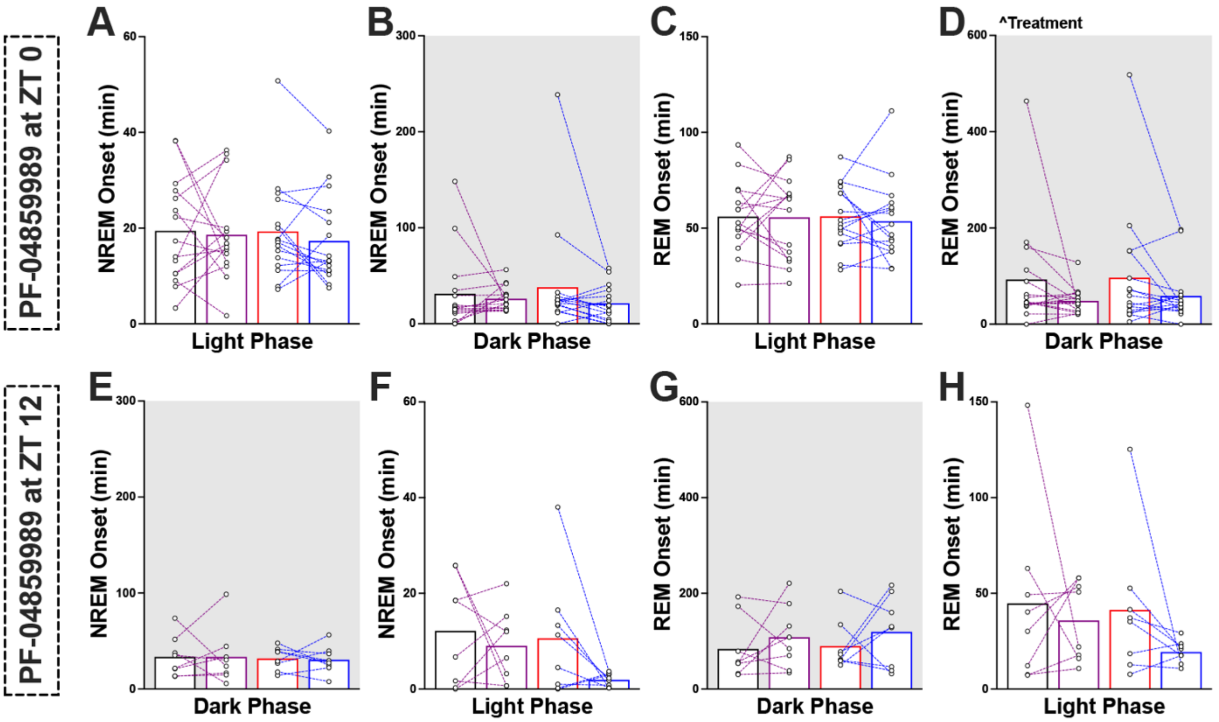


**Supplementary Figure 5. NREM and REM onset following PF-04859989 administration.** Adult ECon and EKyn offspring were treated with vehicle or PF-04859989 (30 mg/kg) at Zeitgeber time (ZT) 0 or ZT 12. **(A)** NREM onset in light phase, sexes combined. **(B)** NREM onset in dark phase, sexes combined. **(C)** REM onset in light phase, sexes combined. **(D)** REM onset in dark phase, sexes combined (Treatment effect $F_{(1, 29)}$= 4.924, P<0.05). **(E)** NREM onset in dark phase, sexes combined. **(F)** NREM onset in light phase, sexes combined. **(G)** REM onset in dark phase, sexes combined. **(H)** REM onset in light phase, sexes combined. Data are mean ± SEM. Two-way RM ANOVA: ^P<0.05. N = 8-16 per group.

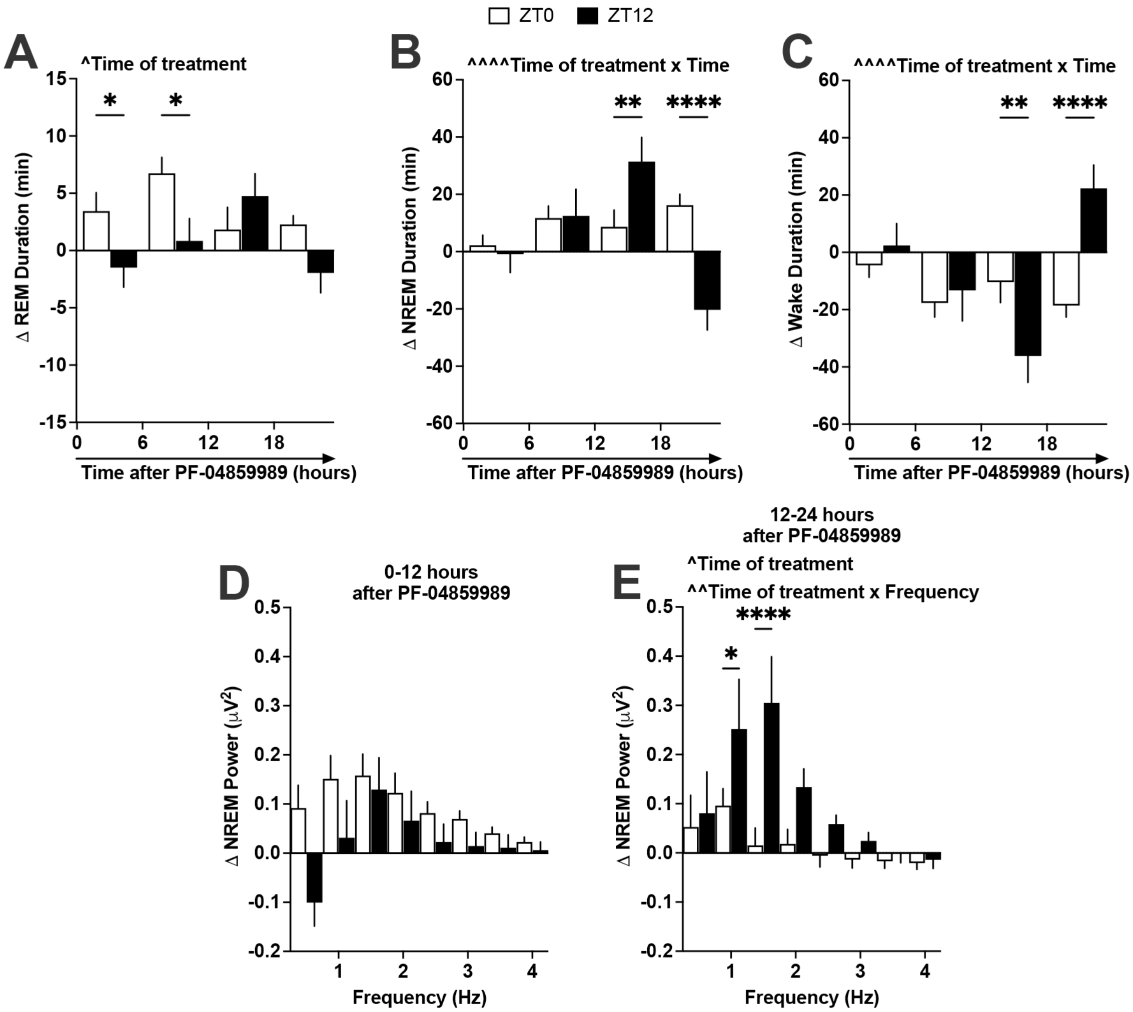


**Supplementary Figure 6. Comparison of changes in sleep architecture between PF-04859989 administration at ZT 0 or ZT 12.** Adult ECon and EKyn offspring were treated with vehicle or PF-04859989 (30 mg/kg) at Zeitgeber time (ZT) 0 or ZT 12. **(A)** 6-hr bins of absolute change in REM duration, sexes and prenatal conditions combined (Time of treatment effect $F_{(1, 175)}$= 5.935, P<0.05). **(B)** 6-hr bins of absolute change in NREM duration, sexes and prenatal conditions combined (Time of treatment x Time interaction $F_{(3, 130)}$= 8.322, P<0.0001). **(C)** 6-hr bins of absolute change in wake duration, sexes and prenatal conditions combined (Time of treatment x Time interaction $F_{(3, 130)}$= 7.712, P<0.0001). **(D)** Absolute change in NREM delta spectral power 0-12 hours after PF-04859989, sexes and prenatal conditions combined. **(E)** Absolute change in NREM delta spectral power 12-24 hours after PF-04859989, sexes and prenatal conditions combined (Time of treatment effect $F_{(1, 45)}$= 6.064, P<0.05, Time of treatment x Frequency interaction $F_{(7, 315)}$= 3.078, P<0.01). Data are mean ± SEM. Two-way RM ANOVA: ^P<0.05, ^^^^P<0.0001 with Fisher's LSD post hoc test: *P<0.05, **P<0.01, ****P<0.0001. N = 16-31 per group.
